# Pneumococcal Colonisation is an Asymptomatic Event in Healthy Adults using an Experimental Human Colonisation Model

**DOI:** 10.1101/652370

**Authors:** Ashleigh Trimble, Victoria Connor, Ryan E Robinson, Carole A Hancock, Duolao Wang, Stephen B Gordon, Daniela M Ferreira, Angela D Wright, Andrea M Collins

## Abstract

**Introduction:** Pneumococcal colonisation is regarded as a pre-requisite for developing pneumococcal disease. In children previous studies have reported colonisation to be a symptomatic event and described a relationship between symptom severity/frequency and colonisation density. The evidence for this in adults is lacking in the literature. This study uses an experimental human pneumococcal challenge model to explore whether pneumococcal colonisation (or co-colonisation with a respiratory virus) is a symptomatic event in healthy adults.

**Methods:** Healthy volunteers aged 18-50 were recruited and inoculated intra-nasally with either *Streptococcus pneumoniae* (serotypes 6B, 23F) or saline as a control. Respiratory viral swabs were obtained prior to inoculation. Nasal and non-nasal symptoms were then assessed using a modified Likert score between 1 (no symptoms) to 7 (cannot function). The rate of symptoms reported between groups was compared and a correlation analysis performed.

**Results:** Data from 54 participants were analysed. 46 were inoculated with *S. pneumoniae* (29 with 6B, 17 with 23F) and 8 received saline. In total, 14 became experimentally colonised (30.4%), all of which were inoculated with 6B serotype. There was no statistically significant difference in nasal (p= 0.45) or non-nasal symptoms (p=0.28) between the pneumococcal inoculation group and the saline group. There was no direct correlation between colonisation density and symptom severity in those who were colonised. In the 22% (12/52) who were co-colonised with pneumococcus and respiratory viruses there was no statistical difference in either nasal or non-nasal symptoms (virus positive p=0.74 and virus negative p=1.0).

**Conclusion:** Pneumococcal colonisation is asymptomatic in healthy adults, regardless of bacterial density or viral co-colonisation.

## Introduction

*Streptococcus pneumoniae* (pneumococcus, SPN) frequently colonises the human nasopharynx, with 40-95% of infants and 10-25% of adults being colonised at any one time(1). Pneumococcal/SPN colonisation rates also vary with geographical location, genetics and socioeconomic background(2). SPN colonisation is a dynamic process. Although multiple pneumococcal serotypes can both simultaneously and sequentially colonise, one serotype is usually the predominant current coloniser(3). In addition interspecies competition occurs between resident flora and potential colonisers including *S.pneumoniae, H.influenza and S.aureus*(4).

Colonisation of the nasopharynx is important as the pre-requisite for SPN infections including pneumonia, sepsis, meningitis and otitis media. Most colonisation episodes will not lead to subsequent disease. Colonisation is also thought to be the predominant source of immunological boosting against SPN infection in both children and adults(5, 6).

SPN colonisation appears to be asymptomatic in murine models(7) and in adults, however the current data are limited(8). Previous studies in children have demonstrated mild nasal symptoms following colonisation(9). Furthermore, a relationship between symptom severity, pneumococcal density and pneumococcal/viral co-colonisation has also been noted in children(10).

Pneumococcal colonisation may cause nasal symptoms in two ways; the bacteria induce host secretions and inflammatory responses or in co-colonised subjects (pneumococcus and virus) due to viral proliferation inducing rhinitis(9). Some studies have also concluded that the presence of respiratory viruses and/or other bacteria within the nasopharynx is the main cause of symptoms; this colonisation in turn increases the rate of pneumococcal colonisation(9).

We have used the novel experimental pneumococcal challenge model (EHPC) to investigate if the process of nasopharyngeal pneumococcal colonisation is symptomatic, causing either nasal symptoms or non-nasal symptoms. This model mimics natural pneumococcal colonisation in healthy human adults and has been used to effectively study mucosal immunity and as a platform to test the efficacy of pneumococcal vaccines in randomised control trials(11).

## Methods

We recruited non-smoking healthy participants aged 18-60 years old. Specimen collection and sample processing were conducted in Liverpool, UK. All participants gave written, informed consent. Ethical permission was granted by local NHS Research and Ethics Committee (REC) (11/NW/0592 Liverpool-East). Exclusion criteria included natural pneumococcal colonisation at baseline, any chronic medical condition or regular medication (study participation could put the volunteer at increased risk of pneumococcal disease) and regular contact with an at-risk individual such as young children (study participation could put the at-risk individual at increased risk of pneumococcal disease).

Participants were nasally inoculated with 8×10^4^, 1.6×10^5^, or 3.2×10^5^ mid-log phase colony forming units (CFU) *S. pneumoniae* (prepared as previously described)(6). Bacterial inoculation density was confirmed by serial dilutions of the inoculation stock onto blood agar (Oxoid). Two serotypes were used; 6B and 23F, both were fully sensitive to penicillin. 46 participants were inoculated with *S. pneumoniae* (SPN) as part of a dose-ranging study and 8 participants inoculated with saline as a control group. Participants were allocated to be inoculated with either 6B, 23F or saline and were blinded to their group.

Pre-inoculation oropharyngeal swabs were assayed for respiratory viruses using multiplex Polymerase Chain reaction (PCR) as previously published (12). The PCR assay panel detected Influenza A and B, Respiratory syncytial virus, Human metapneumovirus, Human rhinovirus, Parainfluenza viruses 1-4 and Coronaviruses OC43, NL63, 229E and HKU1. Nasopharyngeal colonisation was assessed in nasal washes (Nacleiro technique, as previously described) collected at day 2, 7 and 14 post inoculation(13). Pneumococcal colonisation status and density in nasal washes was determined by classical culture as previously described(6, 13).

Participants were prompted to complete a daily symptom log on the day of inoculation (baseline) and daily for 7 days post-inoculation. Symptom log consisted of a 7-point visual analogue scale (a type of Likert scale) which assessed five nasal and five non-nasal symptoms(14). The only modification was removal of ‘mental function’ as a non-nasal symptom (Figure 1). Scores ≥2 were considered ‘symptomatic’. The score awarded at inoculation (day 0) was considered their baseline score, the participant was considered symptomatic if the score went above baseline.

**Figure 1:**
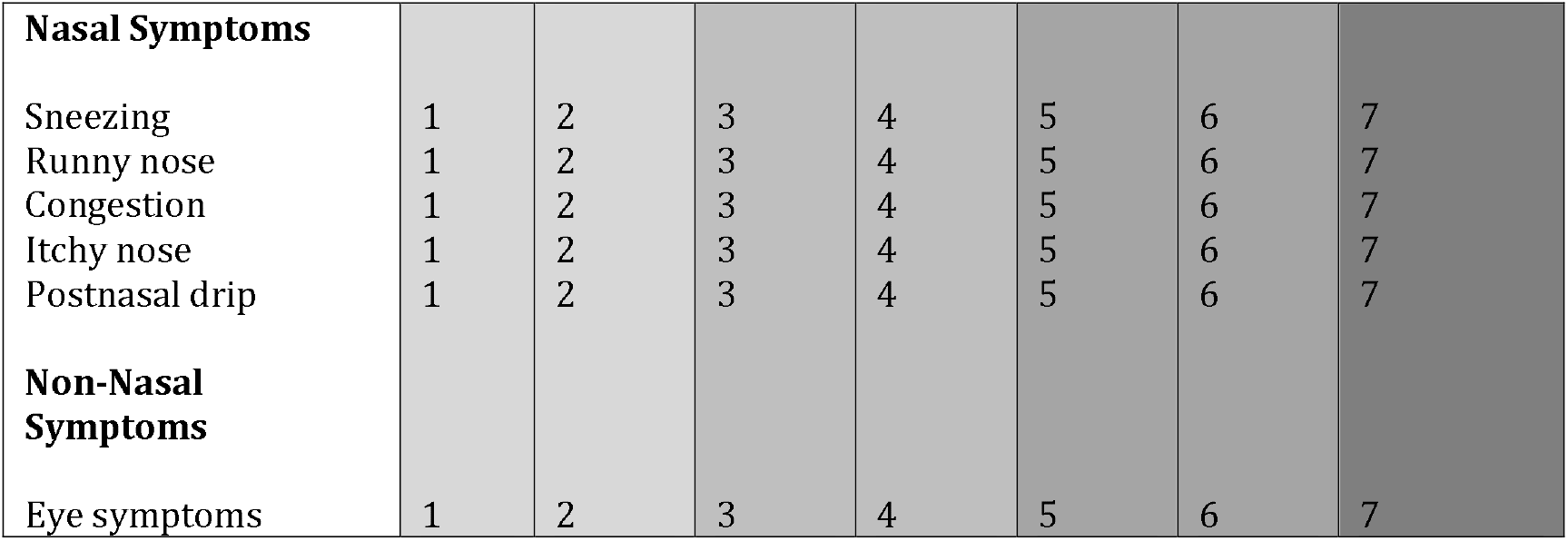
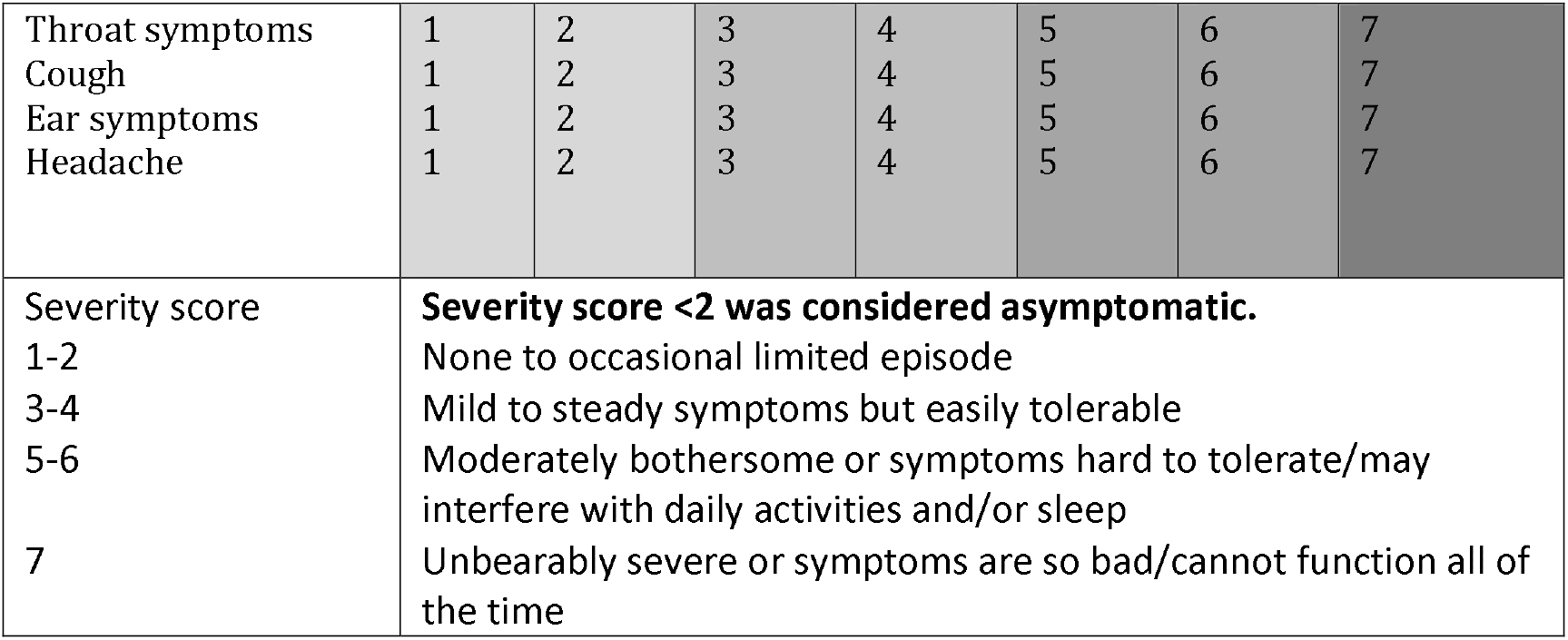
Participant Symptom Log.

Graphical and statistical analyses were performed using GraphPad version 5.0 (GraphPad Software, La Jolla, CA, USA) and Microsoft Excel, with a p-value of <0.05 considered significant. Rates of symptoms reported between groups were compared using Fisher’s exact tests and Chi square where appropriate. Correlation analysis was performed using Spearman’s rank text. The daily symptom logs were collected at the next scheduled visit following completion.

## Results

Fifty-five participants were recruited with an age range of 19-49 years old over a 6-month period from May-October 2014 (check year). Participants with incomplete symptom severity score logs were excluded therefore data from 54 participants were analysed. 46 participants were inoculated with SPN (29 with 6B, 17 with 23F) and 8 with saline (control group). Participants inoculated with 6B, 23F and saline were similar in age and gender distribution. In total, 14 participants became experimentally colonised (30.4%), all of which were inoculated with 6B serotype. None of the participants in the control group developed natural SPN colonisation during the study.

Overall 72% (39/54) of participants reported either or both nasal or non-nasal symptoms during the 7 days post-inoculation. Of these symptoms, similar rates of nasal and non-nasal symptoms were reported. 59% (32/54) of participants reported nasal symptoms and 56% (30/54) reported non-nasal symptoms.

No statistical difference was seen between number of participants who reported symptoms in the experimental SPN positive or negative groups. Similar rates of SPN positive participants reported nasal symptoms (71%, 10/14) and non-nasal symptoms (57%, 8/14) compared to SPN negative participants (50%, 16/32 in nasal and non-nasal). See Figure 2.

**Figure 2.**
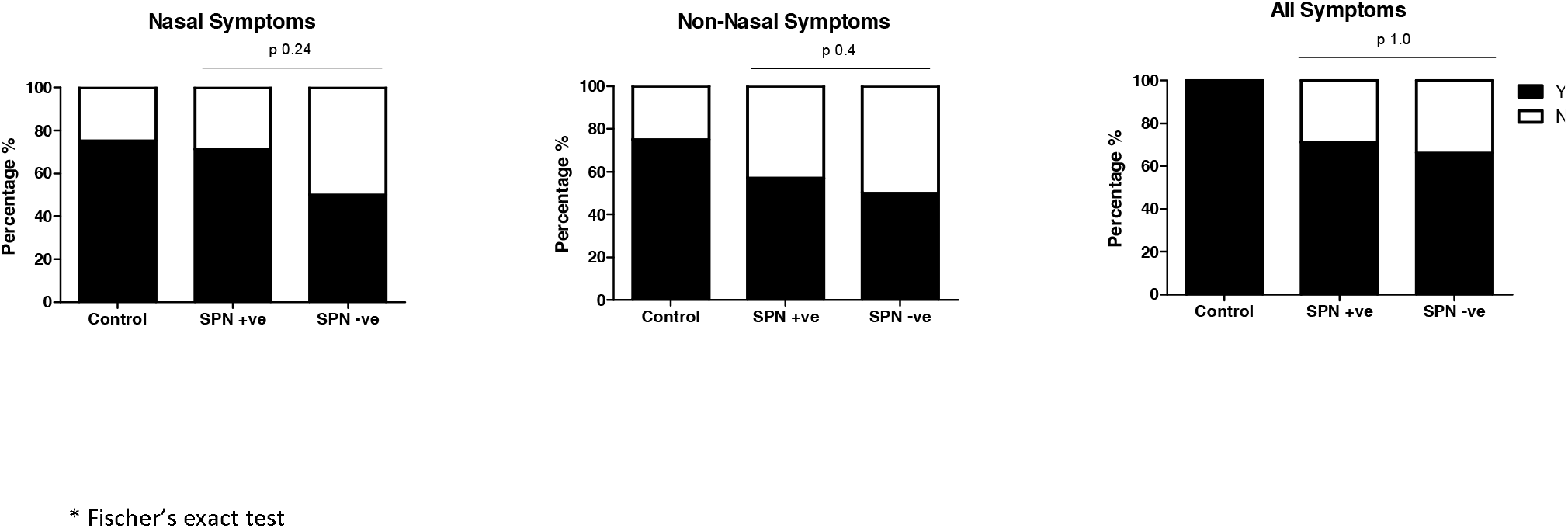
Comparison of nasal and non-nasal symptoms between SPN positive, SPN negative and control participants.

Nasal SPN inoculation did not lead to greater rates of reported symptoms when compared to the saline inoculation group, as show in Figure 3.. Nasal symptoms were reported by 75% of participants inoculated with saline (6/8) compared to 57% (26/46) of those who were inoculated with SPN, no statistical difference was seen (p 0.45). Similarly, no statistical difference was seen with the reporting of non-nasal symptoms 24/46 (52%) post-SPN inoculation compared to post-saline inoculation 6/8 (75%), (p 0.28). Participants that reported ‘any symptom’ were higher in the control group 100% (8/8) compared to 67% (31/46) in the inoculation group, this was not statistically significant (p 0.09).

**Figure 3.**
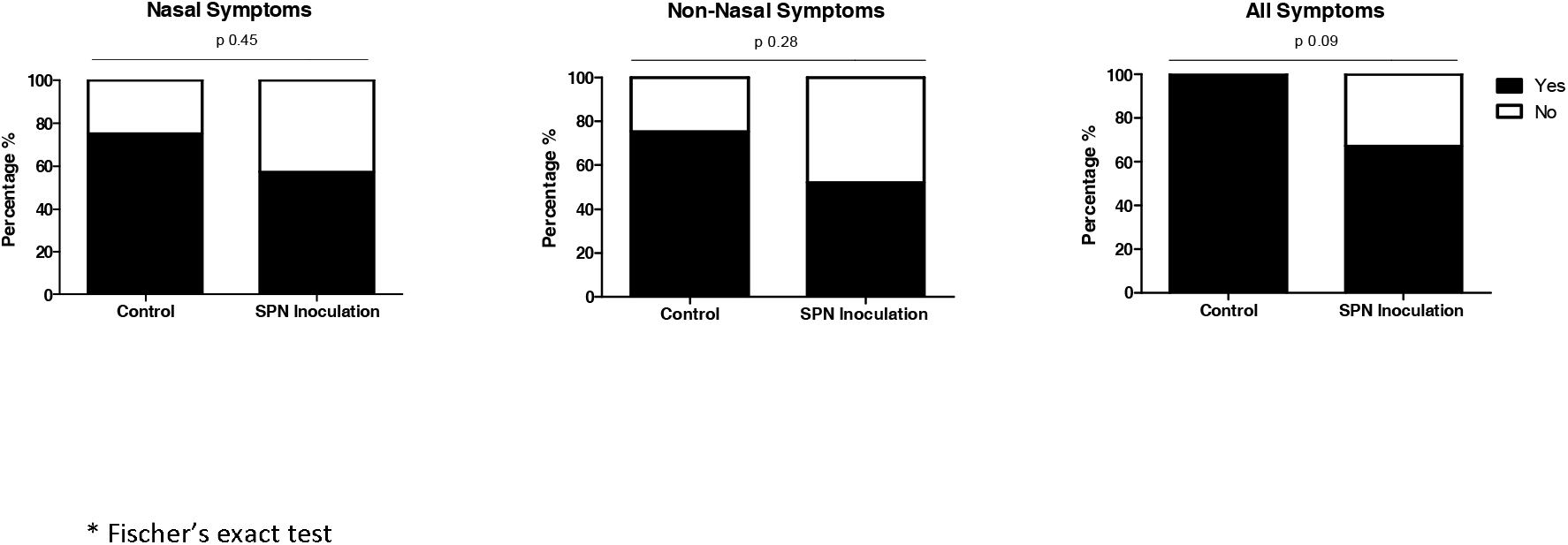
Comparison of nasal and non-nasal symptoms between SPN and saline (control) inoculated groups.

Of the 14 participants colonised with SPN, colonisation density was measured at days 2 and 7. No direct correlation was seen between density and the mean symptom severity score at day 2 and day 7 for nasal and non-nasal symptoms. Figure 4.

**Fig 4:**
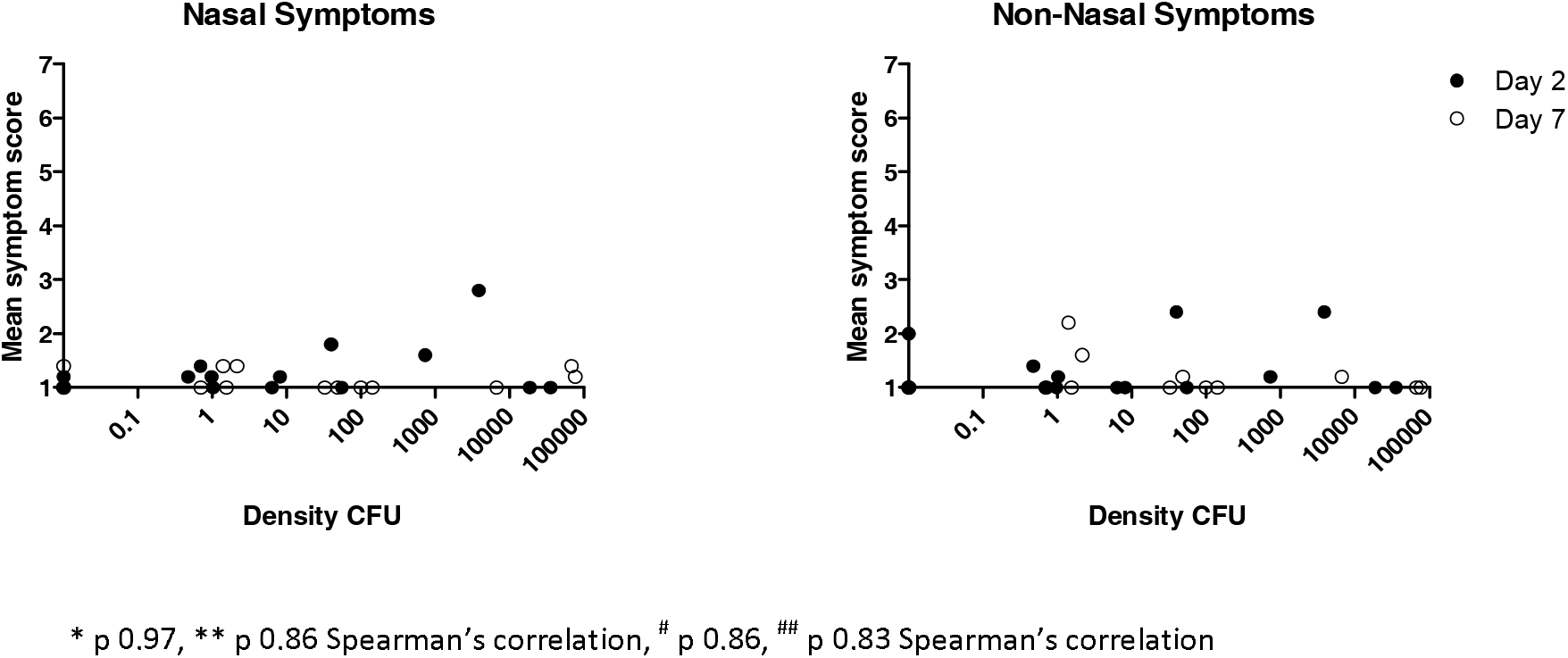
Correlation between pneumococcal colonisation density (SPN positive) and mean nasal severity scores at days 2 and 7.

**Fig 5:**
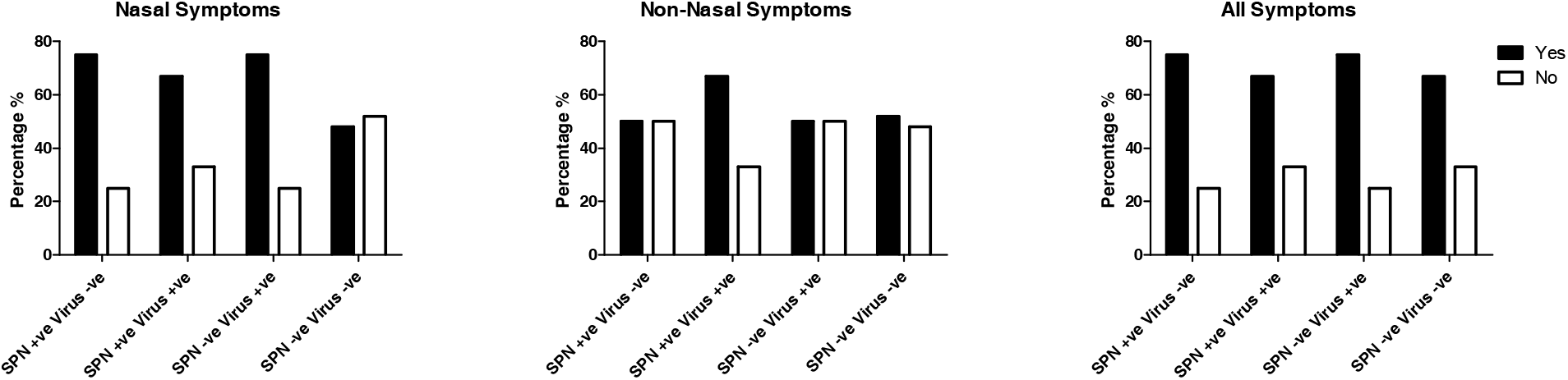
Comparison of nasal, non-nasal and all symptoms between virus and SPN positive and negative participants.

Viral colonisation data was available for 96% (52/54) participants at baseline. Viral colonisation was detected in 22% (12/52) of participants, 2 were inoculated with saline and 10 with SPN [serotype 23F (n=2) and 6B (n=8)].

There was no increase in nasal or non-nasal symptoms in virus positive 8/12 (67%) and 7/12 (58%) respectively compared to virus negative participants 23/40 (58% for both symptoms), p 0.74 and p 1.0.

Experimental SPN colonisation rates were higher in the presence of virus 6/10 (60%) compared to 8/35 (23%) in virus negative participants (p <0.05). Virus and SPN positive participants (Co-colonised) did not report greater rates of nasal or non-nasal symptoms [4/6 (60%) for both symptoms], compared to SPN positive only [6/8 (75%), 4/8 (50%)] and virus positive only [3/4 (75%), 2/4 (50%].

## Discussion

This study shows that pneumococcal (SPN) colonisation in adults is an asymptomatic event. This novel use of a human challenge model allowed for the study of pneumococcal colonisation and symptomology in a controlled environment.

The strengths of this study are the robust methodology used to assess symptom severity(14), the lack of recall bias (due to daily data log completion) and the use of a control group. Using this novel human challenge model, the exact day of pneumococcal inoculation and the onset and termination of each SPN colonisation episode was known allowing association between symptoms and pneumococcal presence and density. The main limitation of our study was the total sample size (n=54).

Although a previous study in adults used a small sample size (n=14) and did not include the methods used to support this conclusion(15), it agrees with our data that pneumococcal colonisation in healthy adults is indeed asymptomatic. Higher symptom severity scores were not a predictor for colonisation.

SPN colonisation is more common in children; therefore, a limitation of this work is the lack of generalisability of results to all age groups, however there is reasonable evidence exists that SPN colonisation in children does cause nasal symptoms(9, 16). One study suggested that the presence of symptoms could be dependent on the serotype of pneumococcus. The authors reported that colonisation with serotype 19F was strongly associated with symptoms such as coryza, sneezing, cough and expectoration. However, these children were recruited from a paediatric hospital emergency room, the study did not report on the diagnosis given to these patients therefore am upper or lower respiratory infection may have been the cause of these symptoms rather than solely colonisation(16).

Rodrigues et al found that rhinitis symptoms, rates of colonisation with SPN and *H. Influenzae (Hi)* in pre-school children decreased with age. Symptoms of rhinitis were reported using the Symptoms of Nasal Outflow Tally (SNOT) score. Both SPN and *Hi* colonisation was strongly associated with increased SNOT scores in children <5 years (p 0.002 and 0.001) whereas colonisation with *S. aureus* was negatively associated with SNOT scores (p 0.04). Interestingly, 40% of asymptomatic children (low SNOT score) were in fact SPN colonised. However, when the data was analysed considering age, the association between SPN colonisation and SNOT scores was weak (p 0.06) whereas the association between SNOT scores and *Hi* colonisation remained strong (p 0.003). They suggest that *Hi* may stimulate rhinitis in children to increase transmission(9). This study does not however report the effect of co-colonisation on symptoms.

Our results suggest that in adults co-colonisation (SPN and virus) is also an asymptomatic process with similar rates of nasal and non-nasal symptoms reported in all groups. Our results did show that asymptomatic viral infection at baseline was associated with the acquisition of SPN colonisation in adults. This is in keeping with results in children which found a virus had a large effect on SPN colonisation even during asymptomatic viral infections(17). They reported that the proportion of children with SPN colonisation was higher during prompted visits for review of URTI symptoms rather than for asymptomatic follow up visits. Due to the small sample size of SPN and virus co-colonisers (n=6), it is difficult to make strong assumptions about the symptomology of this co-infection from our study. Viral swabs were also only performed at baseline (up to 7 days prior to inoculation) therefore we cannot assess correlation between symptoms and viral status at each point, nor was density measured.

In conclusion we have shown that neither nasopharyngeal inoculation nor experimental pneumococcal colonisation cause nasal or non-nasal symptoms in adults. Our results suggest that asymptomatic viral infection prior to nasopharyngeal inoculation or experimental SPN colonisation does not increase nasal or non-nasal symptoms. A better understanding of the process of viral co-infection in adults is needed, further research into the symptoms caused by viral infection prior to or following acquisition of SPN colonisation would add to this study’s preliminary data. A key question, given the difference between adults and children, is the association between colonisation symptoms and transmission; our study confirms that pneumococcal colonisation in adults is asymptomatic, but does not address transmission dynamics.

## Acknowledgements

We would like to thank the members of our DMSC - Professor Robert C Read, University of Southampton (Chair), Professor David Lalloo and Dr Brian Faragher, Liverpool School of Tropical Medicine (LSTM), the respiratory research nurses, Clinical Research Unit staff, the Infectious diseases team, RLBUHT laboratories and clinical trials pharmacy.

This work would not have been possible without members of the LSTM respiratory team; Debbie Jenkins for data input, Jessica Owugha and Shaun Pennington for their involvement with statistical analysis and Jenna Gritzfeld for involvement in bacterial inoculum preparation, sample processing, storage and interpretation of samples and results.

## References

1. Goldblatt D, Hussain M, Andrews N, Ashton L, Virta C, Melegaro A, et al. Antibody responses to nasopharyngeal carriage of Streptococcus pneumoniae in adults: a longitudinal household study. 2005;192(3):387–93.

2. Bogaert D, De Groot R, Hermans PW. Streptococcus pneumoniae colonisation: the key to pneumococcal disease. Lancet Infect Dis. 2004;4(3):144–54.

3. Cobey S, Lipsitch M. Niche and neutral effects of acquired immunity permit coexistence of pneumococcal serotypes. Science. 2012;335(6074):1376–80.

4. Margolis E, Yates A, Levin BR. The ecology of nasal colonization of Streptococcus pneumoniae, Haemophilus influenzae and Staphylococcus aureus: the role of competition and interactions with host’s immune response. BMC microbiology. 2010;10(1):59.

5. Simell B, Auranen K, Kayhty H, Goldblatt D, Dagan R, O’Brien KL. The fundamental link between pneumococcal carriage and disease. Expert Rev Vaccines. 2012;11(7):841–55.

6. Ferreira DM, Neill DR, Bangert M, Gritzfeld JF, Green N, Wright AK, et al. Controlled human infection and rechallenge with Streptococcus pneumoniae reveals the protective efficacy of carriage in healthy adults. Am J Respir Crit Care Med. 2013;187(8):855–64.

7. McCool TL, Weiser JNJI, immunity. Limited role of antibody in clearance of Streptococcus pneumoniae in a murine model of colonization. 2004;72(10):5807–13.

8. Jochems SP, Weiser JN, Malley R, Ferreira DMJPp. The immunological mechanisms that control pneumococcal carriage. 2017;13(12):e1006665.

9. Rodrigues F, Foster D, Nicoli E, Trotter C, Vipond B, Muir P, et al. Relationships between rhinitis symptoms, respiratory viral infections and nasopharyngeal colonization with Streptococcus pneumoniae, Haemophilus influenzae and Staphylococcus aureus in children attending daycare. 2013;32(3):227–32.

10. Fan RR, Howard LM, Griffin MR, Edwards KM, Zhu Y, Williams JV, et al. Nasopharyngeal pneumococcal density and evolution of acute respiratory illnesses in young children, Peru, 2009-2011. 2016;22(11):1996.

11. Collins AM, Wright AD, Mitsi E, Gritzfeld JF, Hancock CA, Pennington SH, et al. First Human Challenge Testing of a Pneumococcal Vaccine - Double Blind Randomised Controlled Trial. Am J Respir Crit Care Med. 2015.

12. Glennie S, Gritzfeld JF, Pennington SH, Garner-Jones M, Coombes N, Hopkins MJ, et al. Modulation of nasopharyngeal innate defenses by viral coinfection predisposes individuals to experimental pneumococcal carriage. Mucosal Immunol. 2016;9(1):56–67.

13. Gritzfeld JF, Wright AD, Collins AM, Pennington SH, Wright AK, Kadioglu A, et al. Experimental human pneumococcal carriage. Journal of visualized experiments: JoVE. 2013(72).

14. Spector SL, Nicklas RA, Chapman JA, Bernstein IL, Berger WE, Blessing-Moore J, et al. Symptom severity assessment of allergic rhinitis: part 1. Annals of Allergy, Asthma & Immunology. 2003;91(2):105–14.

15. McCool TL, Cate TR, Moy G, Weiser JN. The immune response to pneumococcal proteins during experimental human carriage. J Exp Med. 2002;195(3):359–65.

16. Neves FP, Pinto TC, Correa MA, dos Anjos Barreto R, de Souza Gouveia Moreira L, Rodrigues HG, et al. Nasopharyngeal carriage, serotype distribution and antimicrobial resistance of Streptococcus pneumoniae among children from Brazil before the introduction of the 10-valent conjugate vaccine. BMC Infect Dis. 2013;13:318.

17. DeMuri GP, Gern JE, Eickhoff JC, Lynch SV, Wald ER. Dynamics of Bacterial Colonization With Streptococcus pneumoniae, Haemophilus influenzae, and Moraxella catarrhalis During Symptomatic and Asymptomatic Viral Upper Respiratory Tract Infection. Clin Infect Dis. 2018;66(7):1045–53.

